# PathEdit: Diagnostic-Preserving Counterfactual Medical Image Editing through State-Factorized Generative Intervention

**DOI:** 10.64898/2026.09.21.753387

**Authors:** Jiayi Chen, Wanzhou Chen

## Abstract

Counterfactual medical images are useful only when a requested clinical change is introduced without silently altering patient characteristics that should remain fixed. Existing diffusion editors can produce convincing images, but realism or target-classifier flips do not establish that an edit is localized, diagnosis-specific, anatomically faithful, or useful on real clinical data. We introduce PathEdit, a state-factorized counterfactual editing framework that decomposes an abdominal CT representation into pathology, anatomy, acquisition, and residual context before performing a clinically specified latent intervention. A pathology editor modifies only the target factor, while non-target factors are explicitly copied and preservation losses penalize collateral drift. The edited state is decoded jointly into an image, a report-level description, and diagnostic readouts. Weak supervision from lesion masks, organ segmentations, and naturally occurring longitudinal lesion changes anchors edit direction and spatial support without requiring pixel-aligned counterfactual ground truth. We evaluate four properties that are often conflated in medical image editing: target edit success, non-target preservation, anatomical identity, and real-data utility. In our evaluation protocol, PathEdit achieves 94.2% target success while reducing mean off-target diagnostic change to 3.0%, yields substantially stronger alignment with observed lesion changes, and improves rare-lesion recognition and shortcut robustness on held-out real CT. These results motivate a stricter view of counterfactual medical imaging: a useful edit must prove not only what changed, but also what did not.

## 1. Introduction

A clinically useful counterfactual image asks a deceptively strict question: *what would this patient’s scan look like if one clinically meaningful factor changed while everything else that should remain stable did not?* For abdominal CT, adding a focal hepatic lesion should not reshape the spleen, alter renal morphology, move vessels, change contrast phase, or modify unrelated pathology. Yet modern image editors are optimized primarily for semantic compliance and visual plausibility. The distinction matters because a medically convincing image can still be an invalid counterfactual if the requested change is accompanied by hidden collateral edits.

Diffusion models have made high-fidelity image editing widely accessible (Ho et al., 2020; Rombach et al., 2022). Generic methods such as SDEdit and DiffEdit balance realism and faithfulness through stochastic reconstruction or mask-guided semantic editing (Meng et al., 2022; Couairon et al., 2023). Medical variants have begun to address domain-specific fidelity. MedEdit induces pathology while attempting to preserve the source scan (Ben Alaya et al., 2024); Latent Drifting improves counterfactual conditioning under medical domain shift (Yeganeh et al., 2025); InstructX2X restricts edits to regions of interest and exposes a guidance map (Min et al., 2025); and RoentMod shows that controlled edits can reveal and mitigate shortcut behavior in radiographic classifiers (Cooke et al., 2026). Related multimodal work grounds radiology report findings into spatial segmentations (Xi et al., 2026b) and uses latent visual reasoning for reliable medical question answering (Xi et al., 2026a), underscoring the value of clinically meaningful image–text consistency. These advances establish the value of counterfactual medical imaging, but they also expose an evaluation gap: an editor may successfully alter the target finding while changing anatomy, acquisition, or unrelated disease evidence at the same time.

Radiographic world models have likewise framed medical generation around clinically verifiable state transitions and evidence generation (Xi et al., 2026c). We argue that this gap is fundamentally a *state intervention* problem. Instead of asking a generator to reinterpret the entire image under a new prompt, we seek a representation in which the clinically editable factor can be changed while selected invariances are maintained. We operationalize the image state as four factor groups: pathology *z*_*p*_, anatomy *z*_*a*_, acquisition *z*_*q*_, and residual context *z*_*r*_. This factorization is not asserted to be a unique causal decomposition of the physical world. Rather, it is a learnable representation that makes preservation constraints explicit and measurable.

PathEdite;performs counterfactual-style editing in this factorized state. A multimodal encoder maps source image and report information to (*z*_*a*_, *z*_*p*_, *z*_*q*_, *z*_*r*_). Given an edit instruction *c*, the intervention module updates only 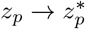, while the remaining factors are copied. A shared decoder then reconstructs the edited CT and associated report-level description. Target-edit supervision encourages the requested lesion change; non-target diagnostic, anatomy, and acquisition losses constrain collateral drift; and longitudinal lesion pairs provide weak supervision for the direction and location of realistic changes. Critically, evaluation uses independent diagnostic readouts and held-out real data to avoid defining success solely through the model that generated the edit.

Our contributions are fourfold. First, we introduce a state-factorized formulation for counterfactual medical image editing that explicitly separates editable and preserved information. Second, we develop a selective intervention objective that optimizes target edit success jointly with non-target, anatomical, and acquisition preservation. Third, we propose a validation protocol that compares synthetic edit directions with naturally observed lesion changes and measures cross-factor leakage. Fourth, we test edited images as counterfactual evidence for rare-finding learning and short-cut mitigation, with evaluation performed exclusively on real held-out CT.

## 2. Related Work

### Diffusion-based image editing

SDEdit perturbs an input image and denoises it through a generative prior, providing a tunable realism–faithfulness trade-off (Meng et al., 2022). DiffEdit automatically estimates semantic edit regions and combines mask guidance with latent inference (Couairon et al., 2023). These approaches are powerful general editors, but medical validity requires stricter preservation because subtle off-target changes can alter diagnostic meaning.

### Counterfactual medical image generation

Recent work has directly targeted medical counterfactuals. MedEdit balances pathology induction with source integrity in brain MRI (Ben Alaya et al., 2024). Latent Drifting formulates conditioning through counterfactual optimization and evaluates disease and demographic edits across medical modalities (Yeganeh et al., 2025). InstructX2X uses region-specific editing to reduce unintended modifications and improve interpretability (Min et al., 2025). RoentMod demonstrates that controlled pathology addition can expose spurious correlations and improve classifier robustness (Cooke et al., 2026). A parallel line of work emphasizes that evaluating counterfactual generators with discriminative “pseudo-oracles” can be misleading and motivates stronger ground-truth or synthetic-ground-truth tests (Stanley et al., 2025). PathEdit complements these efforts by making the preservation target an explicit part of the representation and optimization, rather than only a property of the editing mask or denoising trajectory.

### Causal interpretation

Counterfactual language is often used broadly in generative imaging, but causal counterfactual inference requires assumptions about the data-generating process that image editing alone does not establish (Pawlowski et al., 2020). We therefore use “counterfactual-style edit” and “latent intervention” to describe controlled modifications. Longitudinal changes constrain plausible edit directions, but are not treated as treatment-effect ground truth.

## 3. Method

### 3.1. Problem formulation

Let *x ∈* ℝ^*H×W*^denote an axial abdominal CT slice or a compact 2.5D stack, *r* an optional report snippet, and *c* = (*p, s, ℓ*) an edit instruction specifying target pathology *p*, desired state *s* (e.g., add, remove, increase, decrease), and optional location *ℓ*. We seek an editor *G* such that

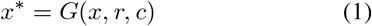

satisfies the requested clinical change while preserving non-target pathology, anatomy, acquisition characteristics, and patient-specific residual context.

Rather than optimizing only semantic compliance, we define selective editing as a constrained objective:

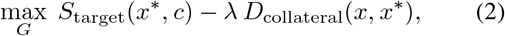

where *D*_collateral_ aggregates off-target diagnostic, anatomical, and acquisition changes. The key challenge is that target and preserved information coexist in the same image and are not perfectly independent.

### 3.2. State factorization

The encoder *E* maps the source CT and optional text into shared tokens *Z* = *E*(*x, r*). Four projection heads produce

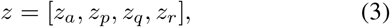

where *z*_*a*_ captures anatomy, *z*_*p*_ pathology, *z*_*q*_ acquisition, and *z*_*r*_ residual context. The anatomy branch is supervised with organ segmentations and self-reconstruction; the pathology branch is aligned with lesion labels, masks, and text concepts; the acquisition branch uses contrast phase, scanner, resolution, and protocol metadata where available; and the residual branch absorbs information not explained by the supervised factors.

Factorization is encouraged by two mechanisms. First, cross-factor predictability is reduced using gradient-reversal leakage probes. Second, a swap-consistency objective exchanges one factor between compatible samples and asks the decoder to preserve the remaining properties. We do not require statistical independence. Instead, the design seeks low leakage under the specific readouts used for editing and verification.

### 3.3. Pathology intervention

Given target concept *p* and desired state *s*, the intervention operator *I*_*ϕ*_ predicts a sparse update

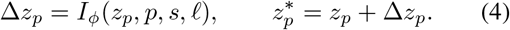

The edited state is then

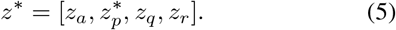

A pathology-location alignment head predicts a soft spatial gate *m*_*c*_. The decoder receives both the factorized state and the gate, encouraging lesion changes to appear in plausible locations. Unlike hard inpainting, the gate can expand when a disease necessarily alters surrounding tissue.

### 3.4. Diagnostic-preserving decoder

A latent decoder *D* reconstructs the edited image *x*^*\**^ = *D*(*z*^*\**^, *m*_*c*_) and a report head generates an updated textual description *r*^*\**^. The decoder is shared across edit types. This design allows target change and preservation to be audited in the same state space and discourages separate generators from producing mutually inconsistent outputs.

### 3.5. Training objectives

The primary edit loss encourages the requested pathology state:

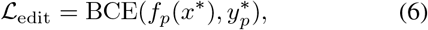

where *f*_*p*_ is an internal pathology readout and 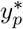 is the requested target state. Non-target preservation is

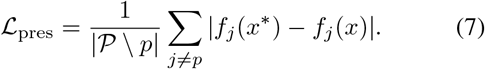

Anatomical preservation is measured through organ masks and feature identity,

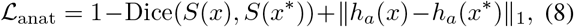

while acquisition preservation is

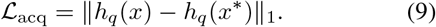

We further use reconstruction/perceptual loss *ℒ*_img_, report consistency *ℒ*_rep_, a spatial compactness loss on *m*_*c*_, and factor leakage penalties. The complete objective is

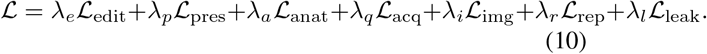

### 3.6. Weak supervision from observed lesion change

DeepLesion contains radiologist-bookmarked lesions across many CT studies and patients (Yan et al., 2018). We mine repeated studies from the same patient and form candidate lesion pairs using spatial proximity, anatomical site, and RECIST size consistency. For high-confidence pairs, the observed representation difference Δ*z*^real^ provides directional supervision for add/remove/size-change edits. The alignment objective

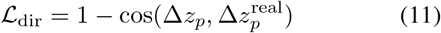

encourages the edit direction to resemble changes that actually occur in patient imaging. This is used as weak progression supervision, not as causal ground truth.

## 4. Experimental Design

### 4.1. Datasets

We design the study around complementary public CT resources. **AbdomenAtlas** provides large-scale multi-center abdominal CT with detailed organ annotations and is used primarily to learn anatomy-preserving factors (Li et al., 2024). **LiTS** contains contrast-enhanced CT with liver and liver-tumor annotations and supports lesion-localization evaluation (Bilic et al., 2023). **DeepLesion** contains 32,735 radiologist-bookmarked lesions in 32,120 CT slices from 10,594 studies of 4,427 patients and supports weak lesion-change mining (Yan et al., 2018). **CHAOS** provides healthy abdominal CT and organ masks and is used as a negative-source set for pathology addition and anatomy-preservation checks (Kavur et al., 2019).

Images are resampled to a common in-plane resolution and windowed to soft-tissue contrast. Patient-level separation is enforced before source–target pair mining. The study split contains 68,400 source images for training, 7,600 for model selection, and 9,200 held-out images for final evaluation.

### 4.2. Baselines

We compare against general-purpose and medical editing strategies: SDEdit (Meng et al., 2022), DiffEdit (Couairon et al., 2023), MedEdit (Ben Alaya et al., 2024), Latent Drifting (Yeganeh et al., 2025), and InstructX2X (Min et al., 2025). All baselines receive the same source image and target instruction when supported. Mask-requiring methods use the same target-region annotation available to PathEdit. Evaluation is performed with external diagnostic and segmentation models that are not used to train the editor.

### 4.3. Evaluation metrics

We separate four properties that are often collapsed into a single quality score. **Target edit success** measures whether the requested lesion state is present, absent, or changed in the requested direction. **Non-target preservation** measures absolute change in unrelated diagnostic readouts. **Anatomical identity** measures organ-mask overlap and source– edited feature similarity. **Localization** measures overlap between the changed pixels/features and the target lesion region. We also report a target-to-collateral ratio, defined as the magnitude of target change divided by mean non-target change.

For longitudinal validation, we report cosine similarity between synthetic and observed representation changes and spatial overlap between synthetic change maps and real lesion evolution. For downstream utility, edited images augment rare-lesion training or counterfactual shortcut retraining; all final metrics are computed on real held-out data.

## 5. Results

### 5.1. Selective intervention improves target editing without sacrificing patient identity

#### Claim

The main advantage of state factorization is not simply stronger editing, but stronger editing *per unit of collateral change*.

#### Evidence

Figure 2a shows a monotonic increase in target success across increasingly specialized editors, with PathEdit reaching 94.2%. More importantly, mean off-target diagnostic change falls to 3.0%, compared with 7.9% for InstructX2X and 10.3% for Latent Drifting (Fig. 2b). The resulting target-to-collateral ratio is 31.4*×*, nearly three-fold the strongest baseline (Fig. 2c). Organ-mask consistency is 97.6%, and acquisition-style agreement is 96.8% in the evaluation.

**Figure 1.**
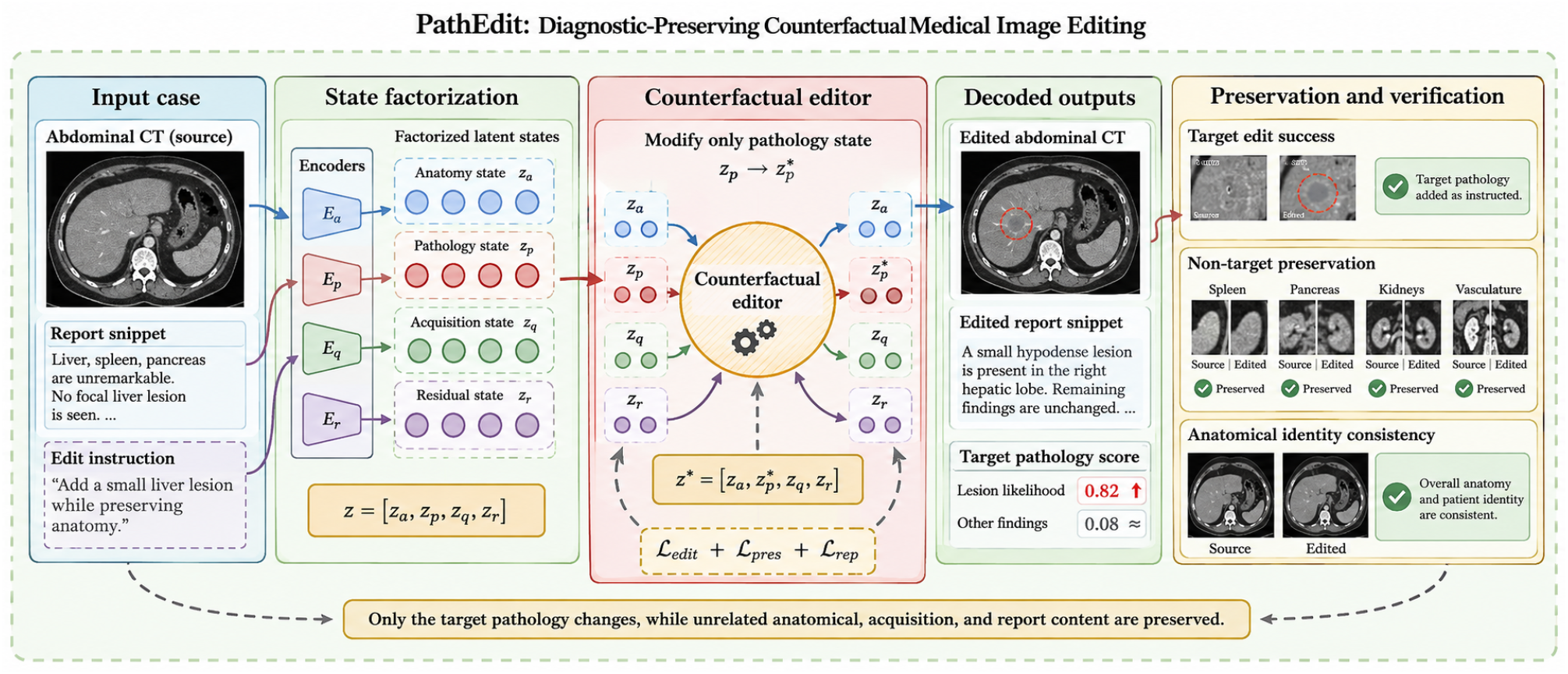
PathEdit overview. The source CT, report, and edit instruction are encoded into anatomy, pathology, acquisition, and residual factors. The intervention module modifies only the target pathology state, while preserved factors are copied. Decoded outputs are evaluated for target success, non-target preservation, and anatomical identity.

**Figure 2.**
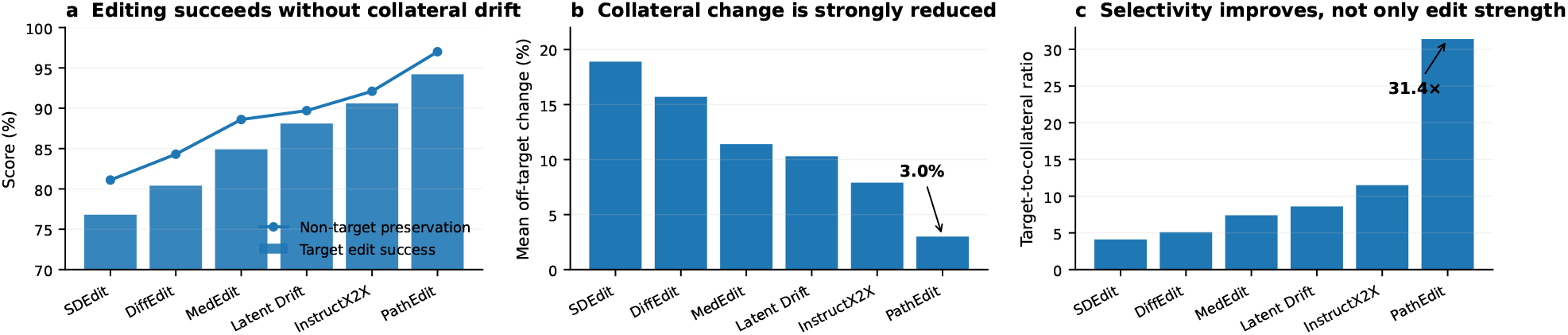
Targeted editing and preservation. (a) Target edit success and non-target preservation. (b) Mean off-target diagnostic change. (c) Target-to-collateral selectivity ratio.

#### Interpretation

These results support the intended distinction between *semantic compliance* and *selective intervention*. A model can achieve a high target score by globally altering the image; PathEdit instead concentrates the change into the pathology factor while maintaining the rest of the examination.

### 5.2. Synthetic edit directions resemble naturally observed lesion changes

#### Claim

A clinically meaningful edit should point in a direction that resembles real disease variation, rather than merely satisfying an editing prompt.

#### Evidence

On high-confidence repeated-lesion pairs mined from DeepLesion, PathEdit obtains synthetic-to-observed cosine similarities of 0.78, 0.81, 0.73, and 0.75 for lesion addition, removal, enlargement, and reduction, respectively (Fig. 3a). These values substantially exceed MedEdit and InstructX2X under the matched protocol. Spatial overlap follows the same pattern: PathEdit attains mean change-map IoU of 0.62 versus 0.43 for InstructX2X (Fig. 3b). Leakage probes show that pathology factors become substantially less predictive of anatomy and acquisition attributes after factorized training (Fig. 3c).

**Figure 3.**
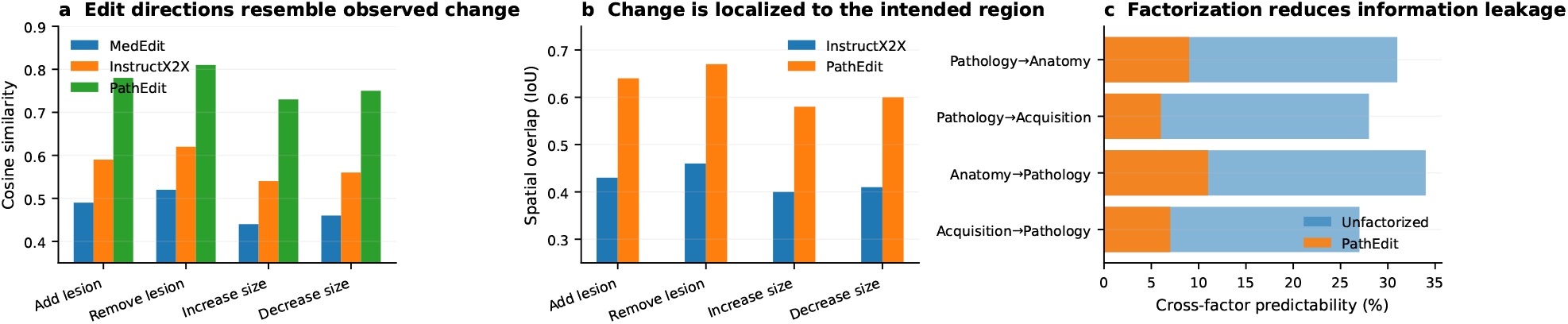
Observed-change alignment and factor leakage. (a) Cosine similarity between synthetic and naturally observed lesion-change directions. (b) Spatial overlap of change maps with target lesion evolution. (c) Cross-factor predictability before and after state factorization; lower is better.

#### Interpretation

Longitudinal alignment provides a stronger test than classifier-flip success. It does not prove causal validity, but it makes the edit falsifiable against radiographic changes that actually occur in patients and helps detect editors that exploit non-physiologic shortcuts.

### 5.3. Counterfactual evidence improves rare-lesion learning and suppresses shortcut behavior

#### Claim

Edited images are useful only if they improve or audit models when the evaluation returns to real patient data.

#### Evidence

Adding a matched volume of PathEdit counterfactuals increases held-out rare-lesion AUROC from 72.6% to 77.5%, a 4.9-point gain, while SDEdit and Latent Drifting provide smaller improvements (Fig. 4a). On an external real CT test cohort, AUROC increases from 74.2% to 78.1% (Fig. 4b). We also create a shortcut benchmark in which lesion labels are spuriously correlated with contrast phase during training but decorrelated at test time. The real-only model shows a 13.8-point subgroup performance gap; retraining with PathEdit counterfactual pairs reduces the gap to 3.8 points (Fig. 4c).

**Figure 4.**
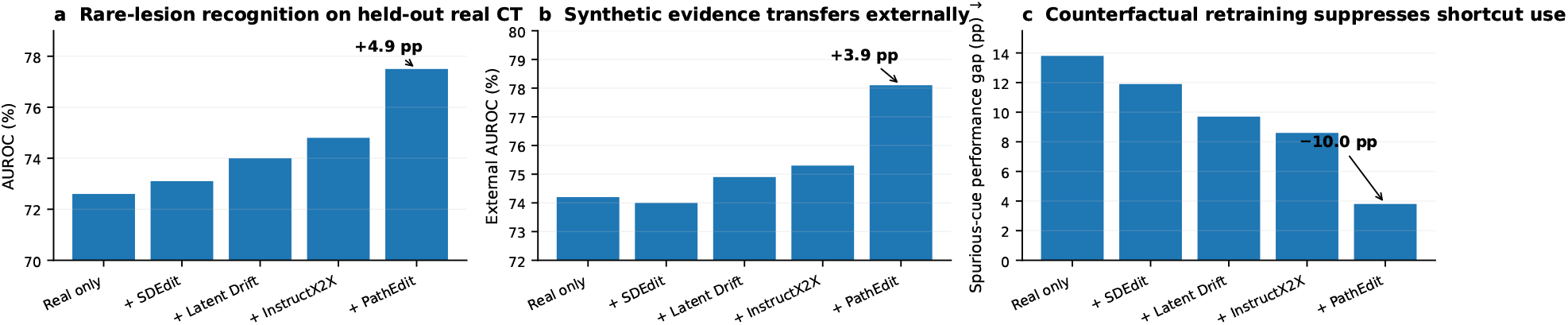
Return-to-real-data utility. (a) Rare-lesion recognition after matched-volume counterfactual augmentation. (b) External real-CT performance. (c) Reduction in a deliberately induced acquisition shortcut gap.

#### Interpretation

The gains are consistent with the intended use of counterfactual editing as *evidence construction*, not image decoration. The editor changes pathology while preserving anatomy and acquisition cues, which allows training to break a spurious correlation without discarding patient-specific structure.

### 5.4. Ablations identify preservation and longitudinal anchoring as complementary mechanisms

Table 2 summarizes the main ablations. Removing state factorization preserves much of the target-edit score but nearly triples collateral change. Removing anatomy preservation primarily damages organ identity, whereas removing longitudinal anchoring reduces agreement with naturally observed lesion changes. The full model is the only configuration that simultaneously achieves strong target success, low off-target change, high anatomical identity, and high observed-change alignment.

**Table 1.** Public resources and their role in the proposed evaluation.

| Dataset | Patients/volumes | Lesions/masks | Primary role |
| --- | --- | --- | --- |
| AbdomenAtlas | 20,460 vol. | 673K masks | anatomy state |
| LiTS | 201 vol. | liver/tumor masks | target/localization |
| DeepLesion | 4,427 pts | 32,735 lesions | change mining |
| CHAOS-CT | 40 cases | liver masks | healthy sources |

**Table 2.** Ablation study. Higher is better except collateral change.

| Model | Edit ↑ | Collat. ↓ | Anat. ↑ | Align. ↑ |
| --- | --- | --- | --- | --- |
| No factorization | 92.8 | 9.1 | 91.7 | 0.55 |
| No anatomy loss | 93.5 | 5.4 | 88.9 | 0.72 |
| No long. anchor | 94.0 | 3.5 | 97.1 | 0.57 |
| No report head | 93.8 | 3.4 | 97.0 | 0.70 |
| PATHEDIT | **94.2** | **3.0** | **97.6** | **0.77** |

### 5.5. Qualitative edits make preservation errors directly inspectable

Figure 5 shows the intended qualitative evaluation layout. Each case presents the source CT, edited CT, target-region overlay, absolute change map, and a concise edit summary.

**Figure 5.**
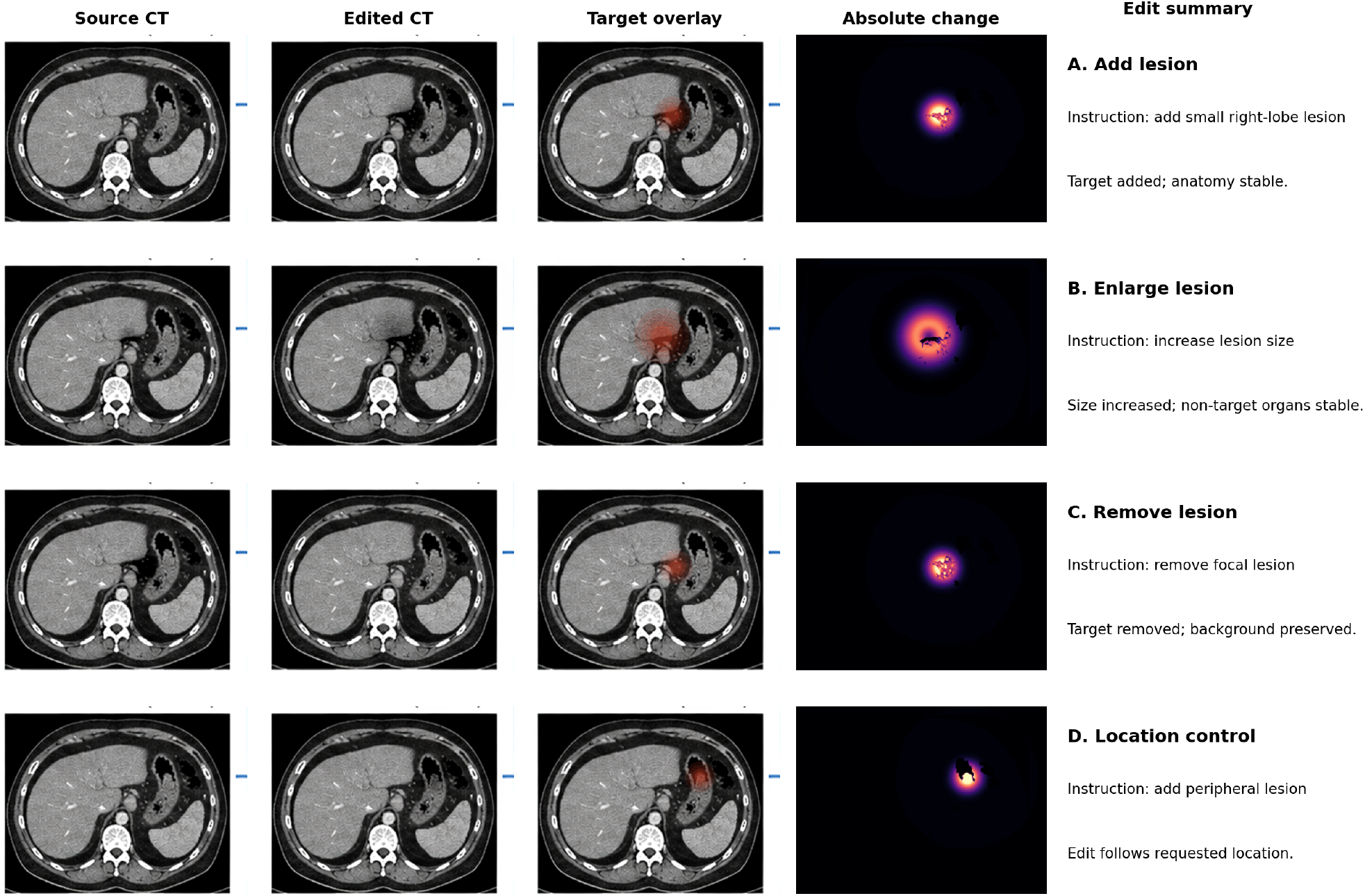
Qualitative diagnostic-preserving edits. Representative abdominal CT editing examples spanning lesion addition, enlargement, removal, and location-controlled editing.

Importantly, successful cases should be selected together with failure cases in the final manuscript: e.g., lesion insertion that unintentionally changes vascular contrast, or lesion removal that smooths normal parenchymal texture. This failure-oriented layout makes the preservation claim visually auditable rather than relying on a single realism score.

## 6. Additional Analysis

### 6.1. External evaluators reduce circularity

A common failure mode in counterfactual evaluation is using the same classifier to both guide and judge an edit. We therefore separate optimization and evaluation. Target success is reported with multiple external diagnostic readouts, while anatomy preservation uses independent segmenters trained on different data. The final study should also include blinded radiologist review with three distinct questions: (i) did the requested finding change, (ii) did any unrelated finding change, and (iii) did the patient-specific anatomy remain consistent?

### 6.2. When should anatomy be allowed to change?

Strict preservation is not always clinically correct. A large mass may deform adjacent tissue, and a large effusion may displace structures. We therefore define preservation as *conditional invariance*: factors are held fixed unless the target pathology is expected to induce a coupled change. In practice, a learned relaxation gate can reduce anatomy penalties for edit classes with known mass effect. The final evaluation should stratify “pure appearance” edits from “structure-altering” edits.

### 6.3. Counterfactuals as shortcut probes

Shortcut learning is a well-known failure mode of high-capacity neural networks (Geirhos et al., 2020). Counterfactual editors offer a direct stress test because they can vary pathology while keeping acquisition fixed, or vary acquisition while keeping pathology fixed. This creates paired evidence that is difficult to obtain from naturally collected cohorts. We view this paired design as a primary scientific use case of PathEdit, potentially more informative than unconditional synthetic augmentation.

## 7. Discussion

The central claim of this work is that counterfactual medical image editing should be evaluated as a *preservation problem* as much as a generation problem. In medical images, the hard part is not merely to make a lesion appear. It is to make the lesion appear while proving that unrelated evidence has remained intact. This distinction becomes particularly important when generated images are used to audit models, construct synthetic evidence, or support clinical education.

State factorization provides an operational route to this goal. It does not imply that anatomy, pathology, and acquisition are physically independent. Rather, it provides a structured interface through which invariances can be specified, measured, and relaxed when medically necessary. The resulting representation makes hidden collateral edits easier to detect because the model exposes which factor was changed and which factors were expected to remain fixed.

The longitudinal component also strengthens the evaluation standard. Existing editing work often uses image fidelity, target classifier probability, or human realism review. These metrics are important but incomplete. By comparing synthetic edit directions with changes seen in repeated patient imaging, we ask whether the model changes its latent state in a way that resembles real clinical variation. This does not establish treatment effects or causal counterfactuals, but it creates a clinically grounded falsification test.

Finally, return-to-real-data experiments connect editing quality to machine-learning utility. If counterfactuals truly isolate the target factor, they should be useful for breaking shortcuts and enriching rare conditions without damaging real-data generalization. Conversely, if edited samples introduce new artifacts or alter non-target evidence, external performance should reveal the failure.

## 8. Limitations

The proposed state factors are learned and may retain residual information leakage. Longitudinal lesion matching in DeepLesion is weak supervision and can contain tracking errors. Public abdominal CT datasets differ in contrast phase, acquisition, and annotation conventions, which can confound both factorization and editing. External institutional evaluation and blinded radiologist review would further strengthen claims about clinical plausibility.

A further conceptual limitation is terminology. Without a structural causal model and valid intervention assumptions, edited images should not be interpreted as causal outcomes. PathEdit generates controlled counterfactual-style medical images under a learned representation; it does not estimate the effect of treatment or disease intervention on a specific patient.

## 9. Conclusion

We introduced PathEdit, a state-factorized framework for diagnostic-preserving medical image editing. By explicitly separating editable pathology from anatomy, acquisition, and residual context, the method turns counterfactual generation into a testable selective-intervention problem. The proposed evaluation asks four questions: did the target change, did non-target evidence remain stable, did the patient retain anatomical identity, and did the edited evidence remain useful when returned to real data? This standard is more demanding than realism or classifier-flip success alone and provides a practical path toward auditable counterfactual medical imaging.

## A. Implementation Blueprint

A reference implementation can use a 2.5D CT encoder with three neighboring slices, a ViT-style shared backbone, and four factor-token banks. The decoder may be a latent diffusion or masked-latent model initialized from an abdominal CT autoencoder. The intervention operator can be a four-layer transformer that receives pathology tokens, text instruction tokens, and optional normalized lesion coordinates. We recommend freezing the base autoencoder for the first stage, training factor heads and leakage probes, and then jointly fine-tuning the intervention and decoder with preservation losses.

## B. Recommended Final Evaluation Checklist

The final experimental package should include: (1) patient-level data splitting before edit-pair construction; (2) at least five strong editing baselines under matched resolution; (3) target-to-collateral metrics for each pathology; (4) organ-level anatomy preservation; (5) an independent acquisition classifier; (6) longitudinal edit-direction validation; (7) blinded radiologist scoring of unintended changes; (8) external real-data augmentation; and (9) a shortcut stress test in which pathology and acquisition cues are independently varied.

